# Malaria entomological surveillance in Kinshasa Province: A longitudinal study

**DOI:** 10.64898/2025.12.22.695884

**Authors:** Fabien Vulu, Melchior M. Kashamuka, Kristin Banek, Ruthly François-Zafka, Pitshou Mampuya, Joseph Atibu, Michael Emch, Victor Mwapasa, Jonathan J. Juliano, Jonathan B. Parr, Rhoel R. Dinglasan, Seth R. Irish, Antoinette K. Tshefu, Thierry L. Bobanga

## Abstract

**Background:** Malaria remains a major public health challenge in the Democratic Republic of Congo. To address the lack of contemporary entomological data, particularly in Kinshasa Province, this four-year study characterized key entomological parameters, including mosquito species composition, abundance, biting behavior, and insecticide resistance.

**Methods:** We conducted a prospective observational cohort study (2018–2022) in three health areas representing rural, peri-urban, and urban settings in Kinshasa Province. Adult *Anopheles* mosquitoes were sampled using CDC miniature light traps and pyrethrum spray catches.

Insecticide susceptibility of adult females was assessed using WHO tube tests, and a generalized estimating equations model evaluated the effects of year, season, site, sampling location, and collection time on mosquito abundance.

**Results:** A total of 1,449 *Anopheles* mosquitoes were collected: 50% in the rural health area of Bu, 49% in peri-urban Kimpoko, and 1% in urban La Voix du Peuple. The species captured were *An. gambiae* s.l. (85%), *An. paludis* (11%), and *An. funestus* (4%). *An. gambiae* s.l. was significantly more abundant indoors and after 10 PM. The overall entomological inoculation rate (EIR) was 4.6 infective bites per person per year, with *An. gambiae* s.l. contributing the highest rate. *An. paludis* showed a localized transmission peak in Impuru during the rainy season (EIR = 8.87). Resistance to DDT and pyrethroids was widespread, with an estimated 0.94 frequency of the kdr L1014F mutation. Bendiocarb and pirimiphos-methyl remained effective.

**Conclusions:** This study confirms that *An. gambiae* s.l. remains the predominant malaria vector and plays a central role in transmission in Kinshasa. However, an emerging role of *An. paludis* in rural settings was observed. These findings underscore the need for vector control strategies tailored to diverse ecological settings. Continued entomological surveillance will be essential to track biting behavior and resistance patterns and ensure that interventions remain effective over time.

**Author summary:** Malaria remains a major health problem in the Democratic Republic of the Congo, but very little recent information is available about the mosquitoes that spread the disease, especially in Kinshasa Province. To help fill this gap, we spent four years studying mosquitoes in rural, peri-urban, and urban communities. We focused on how many mosquitoes were present, which species they were, when and where they bite, and whether they were resistant to the insecticides commonly used in malaria control. We found that most mosquitoes belonged to Anopheles gambiae, the main malaria-transmitting species in Africa. These mosquitoes mostly bit indoors and late at night, when bed nets can offer good protection. However, we also observed mosquitoes biting earlier in the evening and outside, which may reduce the impact of bed nets. Another species, Anopheles paludis, played an important role in malaria transmission in some rural areas, showing that different environments can shape which mosquitoes spread the disease. We also found that many mosquitoes were resistant to widely used insecticides, although some alternative insecticides were still effective. Our findings show why it is important to monitor mosquito behavior and insecticide resistance regularly and to adapt malaria prevention strategies to local conditions.

## Background

Malaria remains a major public health challenge in the Democratic Republic of Congo (DRC), which bears the second highest malaria burden, with an estimated 12.6% of global malaria cases and 11.3% of deaths in 2023 [1]. Kinshasa Province presents a unique setting where various levels of urbanization, landscape changes, and rapid population growth create diverse ecological conditions for malaria transmission. The province contains a mix of rural, peri-urban, and urban environments, each of which may influence vector behavior and malaria transmission dynamics [2]

Fundamental entomological data on malaria transmission remain insufficient, often being outdated or fragmented [1,3–8]. This lack of comprehensive and continuous entomological surveillance hinders efforts to understand local transmission patterns, vector species distribution, insecticide resistance, and the effectiveness of vector control measures. These knowledge gaps make it difficult to effectively target vector control interventions in Kinshasa Province, where the malaria burden varies significantly across different settings [9,10].

Long-lasting insecticidal nets (LLINs) remain the primary malaria prevention tool in Kinshasa and across the DRC [11]. However, LLINs primarily target endophilic *Anopheles* mosquitoes, particularly *Anopheles gambiae* s.l., the main malaria vector in the DRC, which predominantly bites during nighttime hours, with peak activity occurring between 10 pm and 5 am [1,12]. This leaves potential gaps in protection against mosquitoes that bite earlier in the evening, those that bite outdoors or those changing their behavior to avoid LLINs (behavioral adaptations) [13,14]. Moreover, the growing threat of insecticide resistance and changes in vector behavior threaten malaria control efforts in the DRC [6,8].

This study was conducted as part of a broader project integrating epidemiological, parasitological, and entomological approaches to enhance our understanding of malaria transmission across diverse ecological landscapes and time at 7 sites spanning rural, peri-urban, and urban settings in Kinshasa Province [9]. Herein, we characterize and analyze malaria entomological parameters (mosquito abundance, species composition, biting behavior, insecticide susceptibility, and resistance) across varied sites over the course of four years.

## Methods

### Study sites

From 2018 to 2021, we conducted a prospective observational cohort study across seven sites distributed in three health areas (HA) of Kinshasa Province: Bu, Kimpoko, and Voix du Peuple (Fig 1). These sites were purposely selected based on their malaria baseline characteristics [9,15]. Bu and Kimpoko health areas are located in the Maluku 1 health zone, while Voix du Peuple, is in the Lingwala health zone. These health areas represent different environmental settings and levels of malaria transmission: Bu is rural with high transmission, Kimpoko is peri-urban with moderate transmission, and Voix du Peuple is urban with low transmission (Fig 1).

**Fig 1.**
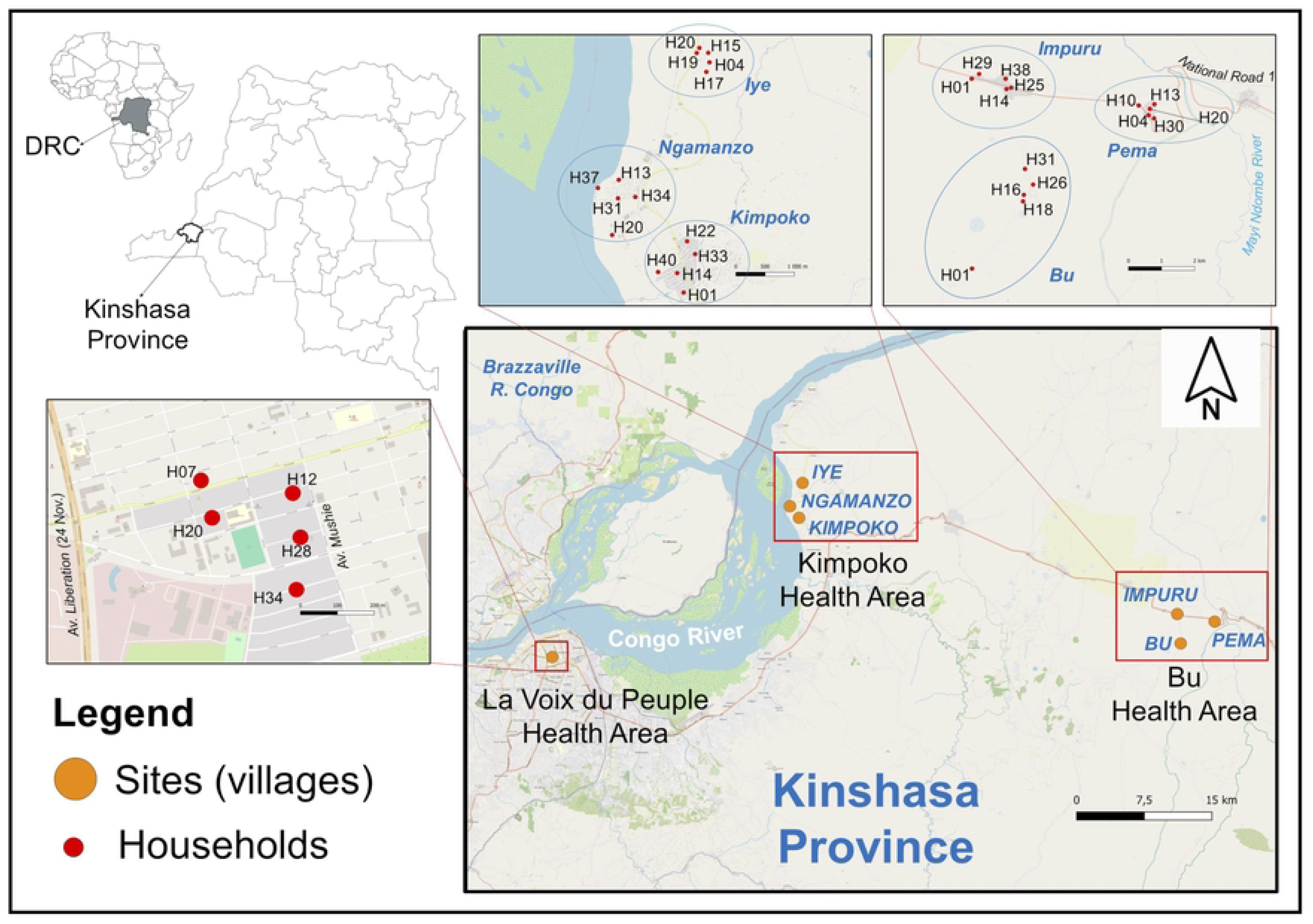
Location of study sites and households. Household locations were deliberately jittered to protect participant confidentiality and to avoid overlapping points on the map. Sites represent distinct settings and transmission levels: Bu (rural, high) on the Bateke Plateau marked by ancient sand dunes, expansive grasslands, patches of wooded savanna, narrow strips of dense gallery forest, and two river valleys; Kimpoko (peri-urban, moderate) within the swampy Maluku 1 corridor along the Congo River; and Voix du Peuple (urban, low) in Lingwala. Within Bu, the Bu site is ∼3 km inland from National Road 1, whereas Impuru and Pema lie along the road. Within Kimpoko, Ngamanzo sits on the riverbank with abundant aquatic vegetation and a small river port, while other sites are farther inland.

Kinshasa Province experiences a tropical humid climate, characterized by a nine-month rainy season from September to May and a three-month dry season from June to August. Mosquito collections occurred in both the rainy and dry seasons. The entomological household sampling methodology has also been previously described [9] In brief, a total of 35 households (five per site) were randomly selected to ensure a spatially representative coverage; 32 households remained enrolled at the last visit.

### Adult mosquito collection and morphological identification

We conducted nine rounds of adult *Anopheles* mosquito collection using U.S. Centers for Disease Control and Prevention Miniature Light Traps (CDC LTs) (John W. Hock Company, model 512) and pyrethrum-spray catches (PSC). Three rounds of mosquito collection took place in both 2018 and 2019, two rounds in 2020, and one round in 2021[9] Due to COVID-19 restrictions in urban areas, the seventh and eighth rounds of mosquito collection were not conducted in La Voix du Peuple HA. However, all sites participated again during the final (ninth round) visit. Two CDC LTs were set per house for three consecutive nights, one indoors (in a sleeping room) and one outdoors (near the entrance). Traps were set at 6 PM, then emptied and reset at 10 PM. The traps set at 10 PM were emptied the following morning at 6 AM. The timing of the trap setting was informed by a survey of children’s and adults’ bedtime, which was found to be around 6-8 PM and 8-10 PM, respectively (unpublished).

After three nights of trapping, PSCs) were performed in these houses using pyrethroid-based commercial insecticide to capture indoor-resting mosquitoes. PSCs were conducted starting at 6 AM, before the opening of household doors and windows, and before the residents begin their daily activities.

*Anopheles* mosquitoes were separated from other mosquito genera based on morphological characteristics and placed individually in 1.5 ml microtubes with silica gel. Samples were then transferred to the insectary of the Department of Tropical Medicine at Kinshasa University for morphological species identification according to Gillies and De Meillon’s keys and stored at – 4°C for future analyses [16,17].

### Larval collection and insecticide bioassays

*Anopheles* aquatic stage collections were conducted on two separate occasions in May 2019 and February 2022 in the Bu and Kimpoko HAs. These periods correspond to the rainy season when the abundance of *Anopheles* larvae was expected to be high. We did not collect *Anopheles* aquatic stages in La Voix du Peuple HA because of the paucity of *Anopheles* mosquitoes in this HA. Upon collection, larvae and pupae were taken to the field insectaries and reared to the adult stage. These insectaries were set up in the primary health centers of Bu and Kimpoko using basic equipment such as larval rearing trays and adult mosquito cages. Temperature and humidity were regularly monitored using digital devices and remained within the required range for all assays.

To assess the insecticide resistance status of *Anopheles* mosquitoes, we conducted WHO insecticide susceptibility tube tests on three- to five-day-old, non-blood-fed F0 adult female *Anopheles* mosquitoes [18]. For each test, four replicates of 20-25 females fed *ad libitum* on a 10% sugar solution were exposed for one hour to insecticide-impregnated papers treated with DDT 4% (Dichlorodiphenyltrichloroethane, Organochlorine), deltamethrin 0.05% (pyrethroid), permethrin 0.75% (pyrethroid), alphacypermethrin 0.05% (pyrethroid), pirimiphos-methyl 0.25% (organophosphate) or bendiocarb 0.1% (carbamate). Controls were also run by exposing two replicates of 20-25 adult female mosquitoes to untreated papers. Both impregnated and control papers were purchased from the Universiti Sains, Malaysia.

### Molecular identification of sibling species of the *Anopheles gambiae* complex

DNA was extracted from the wings and legs of adult-caught *An. gambiae* s.l. following a CTAB (Cetyltrimethylammonium Bromide) technique adapted by the Molecular Biology Unit of the National Institute of Biomedical Research (INRB) (S1 Text). PCRs were conducted using F6 and R5 primers developed by Salako et al. [19], which were adapted from those used by Santolamazza et al. [20] for identifying three species of the *An. gambiae* complex: *An. gambiae* s.s.*, An. coluzzii, and An. arabiensis*, including intermediate individuals (hybrids) (S1 Table).

### Sporozoite rate

To determine the sporozoite rate (SR) of *Anopheles* mosquitoes, we performed enzyme-linked immunosorbent assays (ELISA) on adult female mosquitoes collected in the field. The head–thorax portion of each mosquito was separated and individually homogenized in phosphate-buffered saline containing 0.05% Tween 20 (PBS-T). The homogenates were processed following a modified Wirtz protocol, using the CDC ELISA reagent kit for circumsporozoite protein detection (BEI Resources, NIAID, NIH, USA) [21,22]. Plates were blocked with 1% bovine serum albumin (BSA) in PBS-T, and all incubations were carried out at 37 °C. Peroxidase Substrate Solution B (KPL, 450 mL, 910 Clopper Road, Gaithersburg, MD, USA) and phenol red sodium salt (Sigma-Aldrich) were used in preparing the ABTS substrate (2,2’-Azino-bis (3-ethylbenzothiazoline-6-sulfonic acid)) for color development. Optical density was read at 405 nm after 30 min using an 800 TS microplate reader (BioTek). A sample was considered positive if its OD value was at least twice the mean OD of the negative controls [23].

### Knockdown resistance mutation genotyping

To detect the knockdown resistance (*kdr*) L1014F gene mutation in the sodium channel responsible for the resistance of *Anopheles* mosquitoes to DDT and pyrethroids, DNA was extracted from the wings and legs of mosquitoes as described above and the *KDR*-Li protocol proposed by Huynh et al. was used in adult-caught *Anopheles* specimens [24]. We only sought the *kdr* mutation in *An. gambiae* s.l. because this mutation has only been previously detected in this species [6,25].

### Statistical analysis

Data were analyzed using R version 4.4.1. Statistical analysis was conducted to evaluate the abundance of *Anopheles* species across host-seeking locations (indoor vs. outdoor), time periods, collection sites, HAs, seasons, and years. Chi-square and Fisher’s exact tests were employed to assess associations between categorical factors and mosquito species. For comparisons involving continuous variables, the Mann-Whitney U test was applied. Statistical significance level was set at 0.05.

We evaluated the association between multiple factors (year, season, site, capture location, and time) and the abundance of *Anopheles* spp. using a Generalized Estimating Equations (GEE) regression model. The goal was to account for the correlation between observations within clusters defined by study HA, except for *An. paludis*, for which clustering by HA was not feasible and site-level clustering was used instead. Analyses were conducted using the geepack (version 1.3.12), MASS (version 7.3.60.2), and broom (version 1.0.7) R software packages.

To estimate the association between key factors and mosquito abundance, GEE models were fitted with an exchangeable correlation structure and a Poisson distribution, corrected for overdispersion using the dispersion parameter (θ) derived from a preliminary negative binomial model. Model adequacy was assessed using QIC (Quasi-likelihood under the Independence model Criterion) and Wald tests. The inclusion of covariates resulted in a lower QIC and statistically significant parameter estimates, indicating improved model fit. Missing values (capture location and period time) for PSC collected mosquitoes were handled using weighted frequency imputation, whereby missing values were replaced proportionally based on the observed distribution of each category within the variable.

To calculate the human biting rate (HBR) of *Anopheles* mosquitoes, CDC LTs were used as surrogates for human hosts. While this method can either underestimate or overestimate the actual risk of malaria transmission to humans [26–28], several studies have demonstrated the reliability and validity of using CDC LTs to estimate HBR [29,30]. The HBR was calculated by dividing the number of *Anopheles* mosquitoes captured in a specific area by the number of traps and the number of nights of trapping, yielding the number of bites per person per night. The SR was calculated as the number of mosquitoes testing positive for *Plasmodium* sporozoites by ELISA, over the number of mosquitoes tested, multiplied by 100. The Entomological Inoculation Rate (EIR) was determined by multiplying the HBR by the SR and then by 365, representing the estimated number of infective mosquito bites a person receives per year:

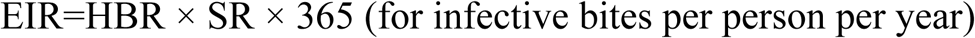

The frequencies of the *kdr* L1014F mutations were calculated using the following formula:

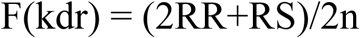

Where RR represents the number of mosquitoes homozygous for the resistant allele, RS represents the number of heterozygous individuals, and n is the total number of tested individuals.

## Results

### *Anopheles* abundance

We collected a total of 1,449 *Anopheles* mosquitoes from participating households, of which 1,288 (88.8%) were captured using CDC LTs. Among these, 937 (72.7%) were collected indoors and 856 (66.4%) after 10 PM. Overall, 1,232 (85.0%) mosquitoes were collected during the rainy season. Nearly all the mosquitoes in this study were captured in the rural site Bu (n = 721, 49.7%) and peri-urban site Kimpoko (n = 716, 49.4%) health areas. By site, the highest abundance of *Anopheles* mosquitoes was recorded in Bu (n = 548, 37.8%) and Kimpoko (n = 531, 36.6%). In contrast, La Voix du Peuple (n = 12, 0.8%) and Pema (n = 37, 2.5%) had the lowest abundance.

### *Anopheles* species

Three *Anopheles* mosquito species were detected: *An. gambiae* s.l. (n= 1,238; 85.4%), *An. paludis* (n=156; 10.7%), and *An. funestus* (n= 55; 3.8%) (Fig 2A). The species distribution varied significantly across the HA (Fisher’s exact test, p<0.001). While 53.3% (n=660) of *An. Gambiae* s.l. were collected in the Kimpoko HA and 46.0% (n=556) in the Bu HA, 86.0% (n=135) of *An. paludis* specimens were found in the Bu HA and 64.0% (n=35) of *An. funestus* were recorded in Kimpoko HA. In the La Voix du Peuple HA, only *An. gambiae* s.l. was recorded (Fig 2B, S2 Table).

**Fig 2.**
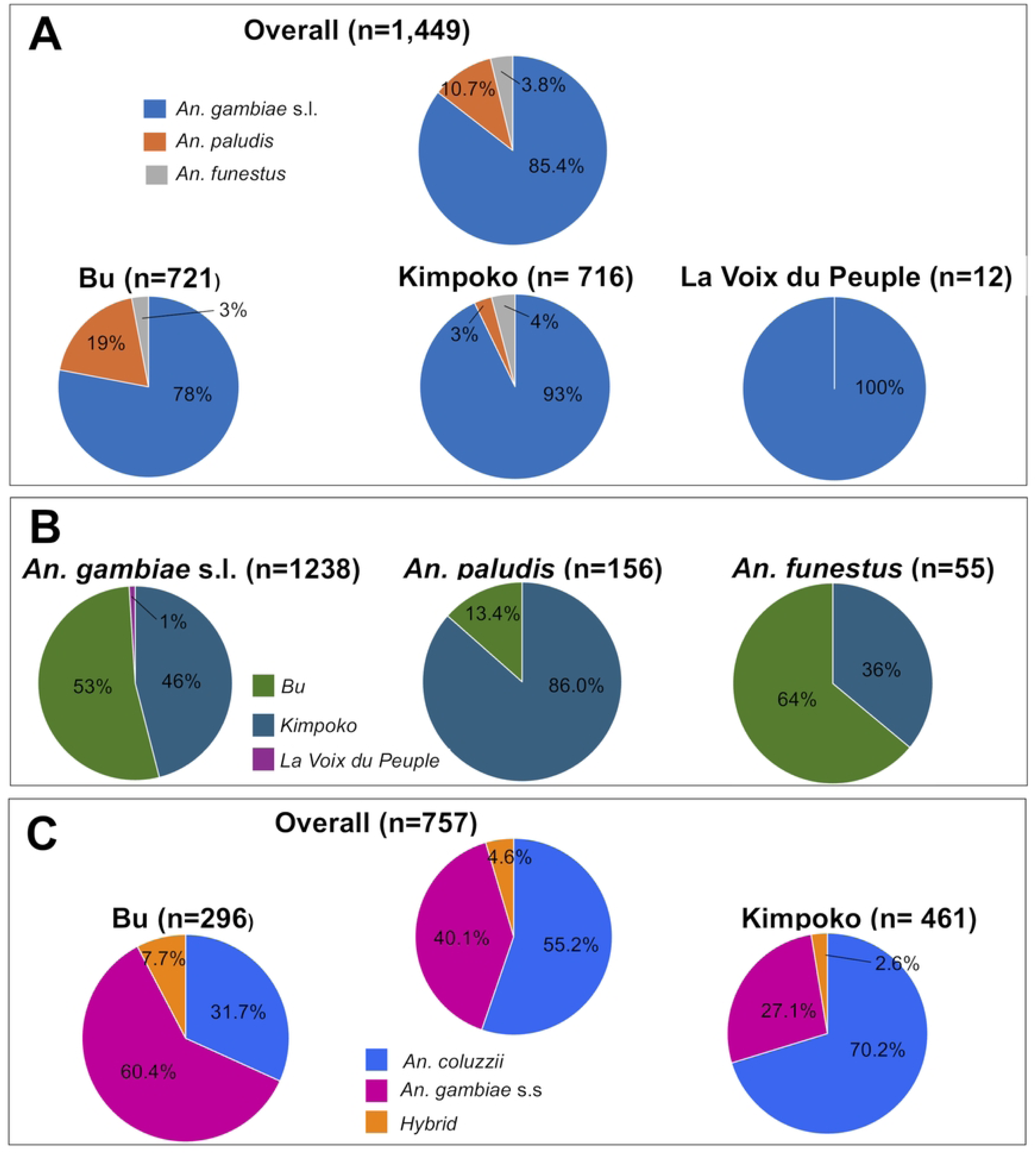
Distribution of *Anopheles* species (A. Abundance of *Anopheles* by health area. B. Distribution of *Anopheles* species across each health area. C. *An. gambiae* s.l. sibling species abundance in each area). Note: Sibling species testing was not performed for La Voix du Peuple due to the very low number (n=12) of *An. gambiae* s.l. specimens collected

The number of *Anopheles* species varied across the different collection rounds, showing notable fluctuations between seasons (Fig 3). Median counts were 141 (*An. gambiae* s.l.), 10 (*An. paludis*), and 8 (*An. funestus*), with significantly higher counts for *An. gambiae* s.l. and *An. paludis* during the rainy season (p = 0.03 and 0.03, respectively). No seasonal difference was observed for *An. funestus* (p = 1.0) (Fig 3).

**Fig 3.**
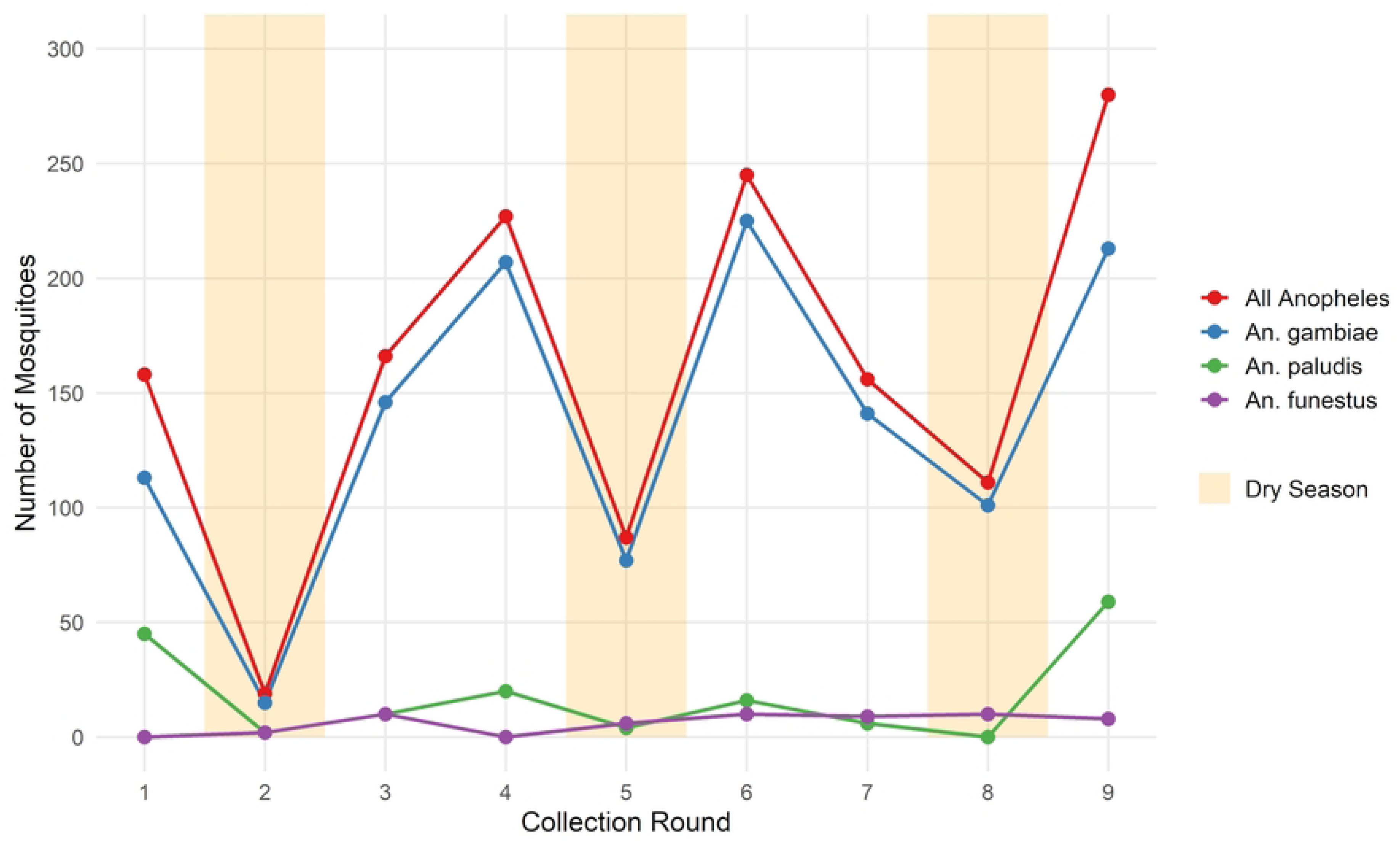
*Anopheles* mosquito species abundance over the study period

Of the 1,238 *An. gambiae* s.l. specimens captured in this study, 757 were molecularly identified to the sibling species level. Among these, 418 (55.2%) were identified as *An. coluzzii*, 304 (40.1%) as *An. gambiae* s.s., and 35 (4.6%) as hybrids of *An. coluzzii* and *An. gambiae* s.s. No *An. arabiensis* was detected. The composition of *An. gambiae* s.l. sibling species varied significantly across HA (χ² test, p < 0.001) (Fig 2C).

### *Anopheles* behavior

Significant differences in mosquito abundance were found by location (indoor vs outdoor) (p < 0.001) and period (before vs after 10 PM) (χ² test, p < 0.001). When analyzed by species, *An. gambiae* s.l. was the prominent species both indoors (p = 0.01) and after 10 PM (p = 0.02). No significant differences in behavior were found for *An. paludis* or *An. funestus* (Table 1).

**Table 1.** *Anopheles* species host-seeking behavior.

| Species | Capture location |  | P | Capture period |  | p |
| --- | --- | --- | --- | --- | --- | --- |
|  | Outdoor | Indoor |  | Before 10 PM | After 10 PM |  |
| *An. gambiae* s.l. | Me: 30 (IQR: 14 - 47)<br>n = 264 (24.3%) | Me: 80 (IQR: 62 - 130)<br>n = 821 (75.6%) | 0.01* | Me: 37 (IQR: 23 - 47)<br>n= 312 (28.7%) | Me: 77 (IQR: 64 - 117)<br>n= 773 (71.2%) | 0.02* |
| *An. paludis* | Me: 4.5 (IQR: 1 – 12.5)<br>n = 68 (44.4%) | Me: 5 (IQR: 2 - 9)<br>n= 85 (55.5%) | 0.68 | Me: 4 (IQR: 2 - 11)<br>n= 98 (64.0%) | Me: 5 (IQR: 1 - 6)<br>n= 55 (35.9%) | 0.72 |
| *An. funestus* | Me: 0 (IQR: 0 - 3)<br>n = 19 (38%) | Me: 3 (IQR: 1 - 6)<br>n= 31 (62%) | 0.27 | Me: 2 (IQR: 1 - 4)<br>n= 22 (44%) | Me: 4 (IQR: 1 – 5)<br>n= 28 (56%) | 0.53 |

Me: Median number of mosquitoes collected per round; IQR: Interquartile range, n= Total number of specimens from all rounds \*Statistically significant (Mann–Whitney U test)

### Factors associated with *Anopheles* abundance

Overall, *Anopheles* abundance was higher in 2019 and 2021, during the rainy season, indoors, and after 10 PM (Table 2). The Bu and Kimpoko sites had significantly higher mosquito abundance than La Voix du Peuple. *An. gambiae* s.l. abundance increased in 2019 and marginally in 2021. It was significantly higher in the rainy season, indoors, after 10 PM, and at all sites except Pema (Table 3). *An. paludis* was significantly less abundant in 2020 but more abundant during the rainy season and at Bu in particular. No association was found with location or collection period (Table 4). *An. funestus* abundance was not associated with most variables, except for a significant increase in 2020 and lower abundance in La Voix du Peuple compared to Iye.

**Table 2.** Factors associated with *Anopheles* abundance.

| Variable | Estimate ($\beta$) | SE | IRR | IRR 95% CI | P |
| --- | --- | --- | --- | --- | --- |
| Year: 2018 | Reference |  |  |  |  |
| Year: 2019 | 0.5 | 0.2 | 1.7 | 1.0 – 2.8 | 0.04* |
| Year: 2020 | 0.1 | 0.3 | 1.1 | 0.6 – 2.0 | 0.61 |
| Year: 2021 | 0.8 | 0.2 | 2.3 | 1.4 – 3.7 | <0.001* |
| Season: Dry | Reference |  |  |  |  |
| Season: Rainy | 0.8 | 0.3 | 2.3 | 1.3 – 4.4 | 0.01* |
| Site: La voix du peuple | Reference |  |  |  |  |
| Site: Bu | 2.2 | 0.3 | 9.5 | 5.2– 17.2 | <0.001* |
| Site: Impuru | 0.8 | 0.4 | 2.3 | 1.1 – 4.9 | 0.03* |
| Site: Iye | 1.0 | 0.3 | 2.9 | 1.5 – 5.5 | <0.001* |
| Site: Kimpoko | 2.4 | 0.3 | 11.3 | 6.4– 19.8 | <0.001* |
| Site: Ngamanzo | 0.9 | 0.2 | 2.4 | 1.5 – 4.1 | 0.001* |
| Site: Pema | 0.7 | 0.4 | 2.1 | 0.9 – 5.0 | 0.08 |
| Time: Before 10 PM | Reference |  |  |  |  |
| Time: After 10 PM | 0.5 | 0.2 | 1.7 | 1.2 – 2.5 | 0.004* |
| Location: Outdoor | Reference |  |  |  |  |
| Location: Indoor | 0.7 | 0.2 | 2.1 | 1.4 – 3.1 | 0.003* |

SE: Standard Error, IRR: Incidence Rate Ratio, CI: Confidence Interval, \*Statistically significant (GEE)

**Table 3.**
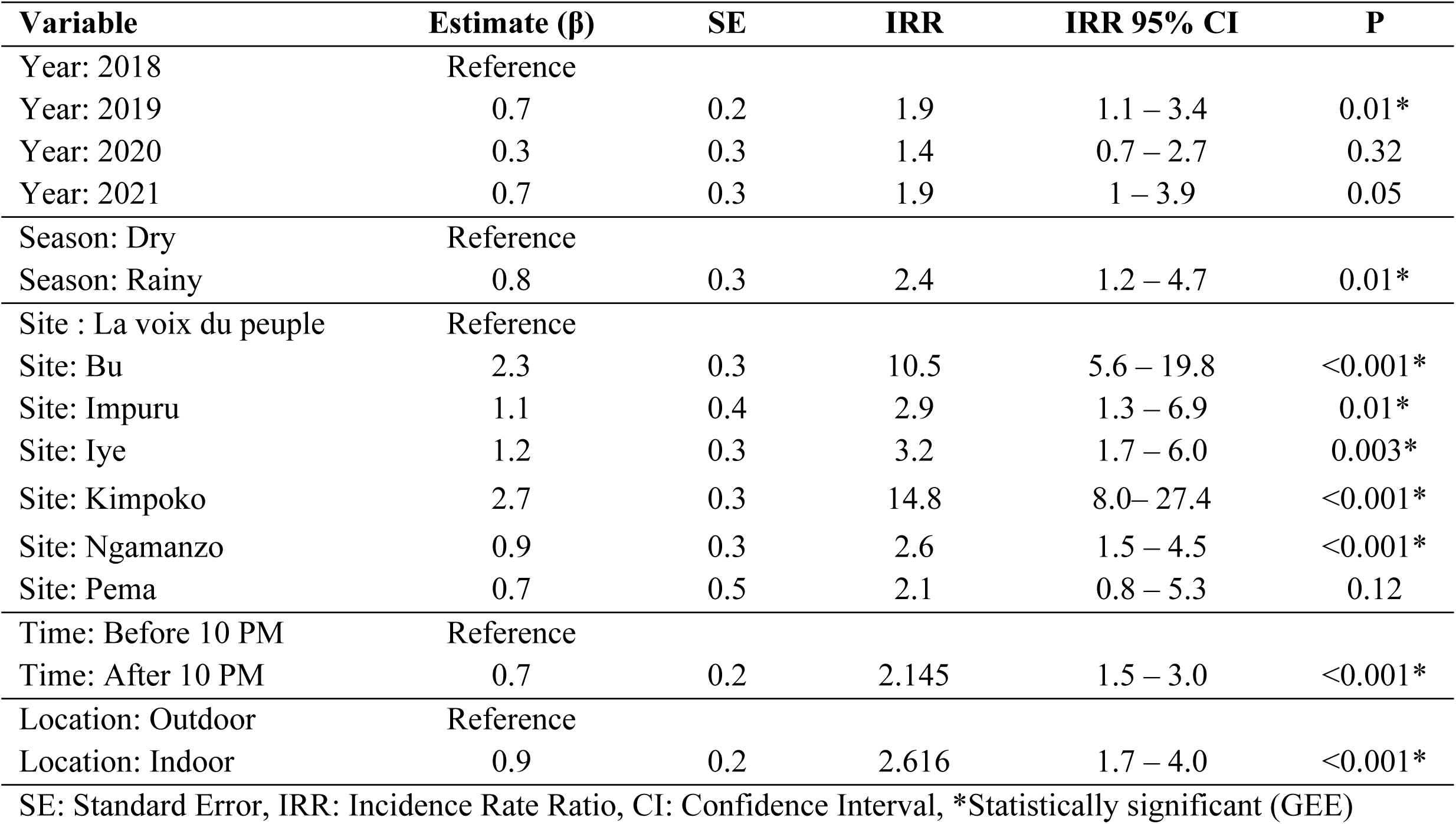
Factors associated with *Anopheles gambiae* s.l. abundance.

| Variable | Estimate (β) | SE | IRR | IRR 95% CI | P |
| --- | --- | --- | --- | --- | --- |
| Year: 2018 | Reference |  |  |  |  |
| Year: 2019 | 0.7 | 0.2 | 1.9 | 1.1 – 3.4 | 0.01* |
| Year: 2020 | 0.3 | 0.3 | 1.4 | 0.7 – 2.7 | 0.32 |
| Year: 2021 | 0.7 | 0.3 | 1.9 | 1 – 3.9 | 0.05 |
| Season: Dry | Reference |  |  |  |  |
| Season: Rainy | 0.8 | 0.3 | 2.4 | 1.2 – 4.7 | 0.01* |
| Site : La voix du peuple | Reference |  |  |  |  |
| Site: Bu | 2.3 | 0.3 | 10.5 | 5.6 – 19.8 | <0.001* |
| Site: Impuru | 1.1 | 0.4 | 2.9 | 1.3 – 6.9 | 0.01* |
| Site: Iye | 1.2 | 0.3 | 3.2 | 1.7 – 6.0 | 0.003* |
| Site: Kimpoko | 2.7 | 0.3 | 14.8 | 8.0– 27.4 | <0.001* |
| Site: Ngamanzo | 0.9 | 0.3 | 2.6 | 1.5 – 4.5 | <0.001* |
| Site: Pema | 0.7 | 0.5 | 2.1 | 0.8 – 5.3 | 0.12 |
| Time: Before 10 PM | Reference |  |  |  |  |
| Time: After 10 PM | 0.7 | 0.2 | 2.145 | 1.5 – 3.0 | <0.001* |
| Location: Outdoor | Reference |  |  |  |  |
| Location: Indoor | 0.9 | 0.2 | 2.616 | 1.7 – 4.0 | <0.001* |

SE: Standard Error, IRR: Incidence Rate Ratio, CI: Confidence Interval, \*Statistically significant (GEE)

**Table 4.** Factors associated with *Anopheles paludis* mosquito abundance.

| Variable | Estimate (β) | SE | IRR | IRR 95% CI | p |
| --- | --- | --- | --- | --- | --- |
| Year: 2018 | Reference |  |  |  |  |
| Year: 2019 | -0.4 | 0.6 | 0.6 | 0.1 – 2.1 | 0.47 |
| Year: 2020 | -1.5 | 0.6 | 0.2 | 0.1 – 0.7 | 0.02* |
| Year: 2021 | 0.8 | 0.6 | 2.2 | 0.7 – 7.2 | 0.19 |
| Season: Dry | Reference |  |  |  |  |
| Season: Rainy | 1.4 | 0.5 | 4.0 | 1.3 – 12.0 | 0.01* |
| Site: Iye | Reference |  |  |  |  |
| Site: Bu | 2.6 | 0.56 | 12.8 | 4.0 – 40.9 | <0.001* |
| Site: Impuru | 1.0 | 0.7 | 2.7 | 0.7 – 10.3 | 0.13 |
| Site: Kimpoko | 0.6 | 0.7 | 1.5 | 0.4 – 6.2 | 0.51 |
| Site: La voix du peuple | -42.2 | 0.7 | 4x10<sup>-19</sup> | 1x10<sup>-19</sup> – 1x10<sup>-18</sup> | <0.001* |
| Site: Ngamanzo | -41.7 | 0.6 | 8x10<sup>-19</sup> | 2x10<sup>-19</sup> – 2x10<sup>-18</sup> | <0.001* |
| Site: Pema | -42.3 | 0.7 | 4x10<sup>-19</sup> | 1x10<sup>-19</sup> – 1x10<sup>-18</sup> | <0.001* |
| Time: Before 10 PM | Reference |  |  |  |  |
| Time: After 10 PM | -0.7 | 0.4 | 0.5 | 0.2– 1.0 | 0.06 |
| Location: Outdoor | Reference |  |  |  |  |
| Location: Indoor | 0.2 | 0.4 | 1.2 | 0.5 – 2.7 | 0.61 |

SE: Standard Error, IRR: Incidence Rate Ratio, CI: Confidence Interval, \*Statistically significant (GEE)

### Human biting rate (HBR), sporozoite rate (SR), and entomological inoculation rate (EIR)

The overall HBR was 0.76 bites per person per night, with the highest rates observed in both Bu (0.83) and Kimpoko (0.81) HAs. *An. gambiae* s.l. had the highest species-specific HBR at 0.64. The overall SR was 1.62% (18/1095 mosquitoes tested), with the highest rates recorded t in both Bu 1.54%, (7/456) and Kimpoko 1.70% (11/648) HAs. The highest site-specific SR was in Pema at 11.1% (1/ 9) (Table 5). The EIR was estimated at 4.49 infective bites per person per year, higher in Bu (4.67) and Kimpoko (5.03), and null in La Voix du Peuple (Table 5). Overall, EIR was higher during the rainy season (5.04 vs 2.51), indoors (14.02 vs 2.66), and after 10 PM (9.27 vs 1.68), except in the Bu HA, where outdoor EIR exceeded indoor. Although *An. gambiae* s.l. had the highest species-specific EIR (3.78), *An. paludis* EIR peaked in Impuru during the rainy season (8.87).

**Table 5.** Entomological risk of malaria transmission.

| Health Areas | Sites | HBR | SR | EIR |  |  |
| --- | --- | --- | --- | --- | --- | --- |
|  |  |  |  | Rainy season | Dry season | All |
| Bu | Bu | 1.77 | 1.13 | 12.05 | 0.00 | 7.30 |
|  | Impuru | 0.53 | 2.13 | 5.40 | 0.00 | 4.12 |
|  | Pema | 0.13 | 11.11 | 8.29 | 0.00 | 5.27 |
|  | **Total** | **0.83** | **1.54** | **10.60** | **0.00** | **4.67** |
| Kimpoko | Kimpoko | 1.81 | 1.59 | 10.11 | 10.38 | 10.57 |
|  | Ngamanzo | 0.24 | 5.26 | 4.29 | 5.23 | 4.61 |
|  | Iye | 0.37 | 0.00 | 0.00 | 0.00 | 0.00 |
|  | **Total** | **0.81** | **1.70** | **4.78** | **4.66** | **5.03** |
| La Voix du Peuple | La Voix du Peuple | 0.05 | 0.00 | 0.00 | 0.00 | 0.00 |
| **Total for all sites** |  | **0.76** | **1.62** | **5.04** | **2.51** | **4.49** |
| ***Anopheles species*** |  |  |  |  |  |  |
| *An. gambiae* s.l. |  | 0.64 | 1.62 | 4.25 | 2.32 | 3.78 |
| *An. paludis* |  | 0.09 | 1.45 | 0.67 | 0.00 | 0.48 |
| *An. funestus* |  | 0.03 | 1.82 | 0.16 | 0.00 | 0.20 |
| **Location** |  |  |  |  |  |  |
| Outdoor |  | 0.41 | 1.26 | 2.66 | 0.00 | 1.89 |
| Indoor |  | 1.11 | 1.86 | 14.02 | 2.99 | 7.54 |
| **Collection Time** |  |  |  |  |  |  |
| Before 10PM |  | 0.51 | 1.46 | 1.68 | 3.78 | 2.72 |
| After 10PM |  | 1.01 | 1.85 | 9.27 | 2.29 | 6.82 |
| **HBR** (Human Biting Rate): Bite per person per night; **SR** (Sporozoite Rate): %; **EIR** (Entomological Inoculation rate): Infective bite per person per year |  |  |  |  |  |  |

### Resistance status to insecticides

In Bu, resistance to DDT was observed in 2019, with a low mortality rates of 25%. By 2022, this mortality rates improved substantially to 89% (Table 6). In 2019, mortality rates for deltamethrin and alphacypermethrin were 86% and 72%, respectively. Deltamethrin mortality remained similar over time, while Alphacypermethrin showed a modest improvement increasing from 72% in 2019 to 82% in 2022. Permethrin, tested only in 2022, showed high resistance with a mortality rate of 55.4%. In contrast, both bendiocarb and pirimiphos-methyl remained fully effective in both years, with 100% mortality. In Kimpoko, strong resistance to DDT was also observed in 2019, but this improved markedly by 2022 (9.7% to 92.7% mortality). Pyrethroid resistance varied: deltamethrin mortality improved from 76.3% in 2019 to 95.8% in 2022, while alphacypermethrin mortality remained stable. Bendiocarb and pirimiphos-methyl were consistently effective across both years, with 100% mortality.

**Table 6.** Insecticide susceptibility status of *Anopheles* mosquitoes in Bu and Kimpoko.

| Insecticide type | Bu health area |  |  |  |  |  | Kimpoko health area |  |  |  |  |  |
| --- | --- | --- | --- | --- | --- | --- | --- | --- | --- | --- | --- | --- |
|  | 2019 |  |  | 2022 |  |  | 2019 |  |  | 2022 |  |  |
|  | n | Mortality | Status | n | Mortality | Status | n | Mortality | Status | n | Mortality | Status |
| DDT 4% | 80 | 25% | R | 100 | 89% | R | 80 | 9.7% | R | 100 | 92.7% | SR |
| Permethrin 0.75% | 80 | 55.4% | R | 100 | – | – | 80 | 54.2% | R | – | – | – |
| Deltamethrin 0.05% | 80 | 86 % | R | 100 | 86.3% | R | 80 | 76.3% | R | 100 | 95.8% | SR |
| Alphacypermethrin 0.05% | 80 | 72% | R | 100 | 82.8% | R | 80 | 87.8% | R | 100 | 88.7% | R |
| Bendiocarb 0.1% | 80 | 100% | S | 100 | 100% | S | 80 | 100% | S | 100 | 100% | S |
| Pirimiphos-methyl | – | – | – | 100 | 100% | S | – | – | – | 100 | 100% | S |

n = number of tested mosquitoes, S: Susceptible, SR: Suspected resistant, R: Resistant

### Markers of insecticide resistance (*kdr* mutation)

Of the 709 *An. gambiae* s.l. specimens successfully genotyped for the *kdr* mutation, 259 were from Bu, 448 from Kimpoko, and two from La Voix du Peuple. The overall frequency of the L1014F mutation was high (0.944), with slightly lower values in Bu (0.918) than in Kimpoko (0.959) (p = 0.04). Mutation frequencies were similar across sibling species; *An. coluzzii* (0.949), *An. gambiae* s.s. (0.940), and hybrids (0.913) (p = 0.81). The majority of specimens (n = 662; 93.3%) were homozygous for the resistant allele.

## Discussion

This study reveals three key findings with direct implications for malaria control in Kinshasa Province. First, *An. gambiae* s.l. remains the dominant malaria vector, exhibiting high indoor and late-night biting activity, which supports the continued use of long-lasting insecticidal nets (LLINs). However, the data also reveal specific gaps in protection, as some *An. gambiae* s.l. subpopulations exhibited early-evening biting (before 10 PM) and outdoor biting behavior, which reduce the effectiveness of LLINs that mainly protect people sleeping indoors late at night. Second, malaria transmission risk varied geographically, but the similarity in entomological inoculation rates (EIRs) between rural (Bu) and peri-urban (Kimpoko) areas indicates that substantial transmission persists in both settings. This suggests that interventions should not only be geographically tailored but also maintained at scale across both rural and peri-urban zones.

Third, widespread resistance to pyrethroids and near-fixation of the L1014F kdr mutation threaten the efficacy of current LLINs, although the preserved susceptibility to bendiocarb and pirimiphos-methyl provides valuable alternatives for complementary control tools. Collectively, these findings underscore the importance of sustaining and strengthening malaria control efforts through the continued use of LLIN-based strategies, complemented by resistance-informed insecticide selection and adaptive approaches that account for local ecological differences, supported by ongoing entomological surveillance to ensure long-term effectiveness.

The predominance of *An. gambiae* s.l. across study sites confirms its central role as the primary malaria vector in Kinshasa, consistent with prior findings in both the province and the DRC [1,4,31–35]. Its strong preference for indoor and late-night biting supports current LLIN-based strategies [1,8,11,36–38]. However, although the proportion was low, the occurrence of bites outdoors and before 10 PM suggests early and outdoor biting behavior, potentially allowing for residual transmission despite LLIN coverage. This vector behavior has been reported in other African settings [39]. At the same time, *An. paludis* accounted for approximately 11% of the total Anopheles population and contributed about 10% of the overall EIR, with a localized peak in Impuru during the rainy season (EIR = 8.87), highlighting its potential role in seasonal transmission. The presence of other anopheline species, particularly *An. paludis*, which significantly contributed to the local EIR at the Impuru site, supports the importance of context-specific vector surveillance and control [40]. Although often considered a secondary vector, *An. paludis* is known to exhibit more exophagic and early-evening biting tendencies compared with *An. gambiae* s.l., behaviors that can facilitate outdoor or pre-bedtime transmission and reduce the effectiveness of LLINs. In such settings, conventional tools like LLINs may be insufficient, and additional interventions such as outdoor residual spraying (IRS) and insecticide-impregnated screens should be considered to address early and outdoor biting mosquitoes.

We identified two sibling species of the *An. gambiae* complex, *An. coluzzii* (55%) and *An. gambiae* s.s. (40%), but not *An. arabiensis*, which is typically restricted to eastern DRC [11,41–43]. This species distribution aligns with previous reports in Kinshasa [6,35,37,44–46]*. An. coluzzii* was predominant in Kimpoko, likely due to its opportunistic breeding behavior and preference for more stable, semi-permanent water bodies, conditions favored by the proximity to the Congo River and the presence of marshlands, fishponds, and slow-flowing streams. In particular, the Ngamanzo site, located directly along the riverbank and characterized by abundant aquatic vegetation and a small river port, provides suitable habitats for *An. coluzzii*, which is known to tolerate moderate organic pollution and is well-adapted to anthropogenic environments [47,48]. In contrast, *An. gambiae* s.s. dominated in Bu, a rural area on the Bateke Plateau, where breeding sites are primarily temporary, rain-fed, and less disturbed, conditions that align with its preference for small, clean, sunlit water bodies with minimal vegetation and low organic content. These observations are consistent with previous findings that highlight the ecological divergence between these two sibling species [47,49–51]. Low hybridization rates were detected between the two sibling species, consistent with earlier findings [45].

The high malaria transmission risk observed in Kimpoko and Bu is likely driven by favorable ecological conditions that support elevated mosquito densities [15,52,53]. In Kimpoko, environmental conditions appear to sustain relatively stable transmission across both seasons. In contrast, Bu shows marked seasonal variation, with increased transmission risk during the rainy season, likely due to the temporary availability of suitable breeding habitats following rainfall. The absence of transmission in the urban La Voix du Peuple HA likely reflects the effects of urbanization, including reduced breeding habitats. These patterns are consistent with recent studies reporting low malaria prevalence in urban areas of Kinshasa and higher prevalence in peri-urban and rural areas [10,53,54].

Insecticide resistance adds another layer of complexity to vector-control efforts in the study area and across the DRC. Consistent with national trends, our findings reveal high resistance to pyrethroids, and the near fixation of the L1014F *kdr* mutation in both Bu and Kimpoko, posing a significant threat to LLIN efficacy and contribute to ongoing transmission despite high net coverage [6,7,35,37,45]. Yet, the confirmed susceptibility to bendiocarb and pirimiphos-methyl in the same mosquito populations provides viable alternatives to pyrethroids. Interestingly, we observed a marked improvement in mosquito mortality to DDT between 2019 and 2022. This unexpected trend is intriguing, especially considering that *Anopheles* population has been highly resistant to DDT in most of the studies in DRC [6,7,12,45]. A possible explanation is that DDT has not been in active use in routine vector control or agriculture for many years, thereby reducing the selective pressure that initially drove resistance. It raises important questions about the drivers and dynamics of resistance, which future studies should investigate.

The findings of this study have important implications for malaria control in the DRC, particularly in Kinshasa. The observed variations in mosquito abundance, species composition, and transmission risk across different ecological settings highlight the complexity of local transmission dynamics. While results support the continued use of LLINs as the cornerstone of malaria prevention, vigilance is needed to detect shifts in mosquito behavior that could influence intervention effectiveness.

The identification of *An. paludis* as a potential primary vector in specific sites further emphasizes the need to expand entomological surveillance to detect emerging vector species and guide timely adjustments to intervention strategies. Moreover, the detection of insecticide resistance, particularly to pyrethroids, underscores the importance of regular tracking to inform insecticide choice. Altogether, these insights emphasize the need for adaptive, locally informed vector control approaches in the DRC.

### Strengths and limitations

This study is one of the first to measure entomological indicators within the context of an ongoing cohort study, but several limitations exist. First, funding constraints limited sampling to five households per study site per round, limiting our ability to detect household-level variations. Second, using the CDC LTs as proxies for human biting rates may have introduced measurement biases. Moreover, *kdr* mutation testing was not performed after insecticide bioassays, preventing direct assessment of phenotypic-genotypic resistance linkage. Additionally, larval and household sampling were not always aligned. Although collections occurred in both wet and dry seasons, calculating annual EIR by multiplying the daily EIR by 365 may overestimate transmission by not fully accounting for seasonal variation. Continuous year-round sampling would improve accuracy.

At the time of testing, the correct dose of pirimiphos-methyl (0.25%) was used, however, WHO updated the recommended dose in 2022. Future susceptibility testing should follow the revised protocol to ensure comparability and accuracy. Chlorfenapyr susceptibility was not evaluated, an important gap given the increased deployment of chlorfenapyr-based LLINs in the DRC. Despite these limitations, the study provides valuable data on mosquito abundance, species composition, behavior, and resistance to insecticides to inform vector control strategies in the DRC.

## Conclusions

This entomological study underscores the importance of adaptive and locally informed malaria control strategies in Kinshasa Province. The dominance of *An. gambiae* s.l., its predominantly indoor and late-night biting behavior, and the high EIRs in rural and peri-urban areas reinforce the continued relevance of LLIN-based interventions, while also exposing critical gaps in protection due to subpopulations of *An. gambiae* s.l. exhibiting early and outdoor biting. The emergence of *An. paludis* as a potential vector in specific ecological niches and the widespread resistance to pyrethroids complicate the malaria transmission and control landscape. Preserved susceptibility to bendiocarb and pirimiphos-methyl offers promising alternatives for vector control. Tailoring interventions to local resistance patterns and ecological contexts, via integrated vector management by combining complementary tools such as indoor residual spraying, larval source management, and spatial repellents, coupled with sustained entomological surveillance, remains essential to ensure that malaria control efforts remain effective, equitable, and responsive to the diverse ecological realities in the DRC.

## Acknowledgements

The authors are grateful to all individuals and institutions who contributed to this study. We wish to express our sincere appreciation to the entomology field team, composed primarily of community health workers, for their dedication and tireless efforts in mosquito collection and processing. We also thank all IRIDA cohort study field team for their support and partnership in the field. Finally, we greatly appreciate the commitment and collaboration from study participants. We acknowledge with deep respect the memory of Dr. Steven R. Meshnick, whose visionary contributions to malaria research and mentorship continue to inspire our work.

The authors used an artificial intelligence language model for English language editing during the writing process. However, the manuscript is original to the authors, who take responsibility for its content.

## Supporting information

**S1 Text. DNA extraction protocol.**

(DOCX)

**S1 Table. Primers used in the study.**

(DOCX)

**S2 Table. Anopheles abundance across sites.**

(XLSX)

## Notes

### Competing Interest Statement

The authors have declared no competing interest.

## References

1. World Health Organization. World malaria report 2023. Geneva: WHO; 2023.

2. Kang YO, Yabar H, Mizunoya T, Higano Y. Optimal landfill site selection using ArcGIS multi-criteria decision-making (MCDM) and analytic hierarchy process (AHP) for Kinshasa City. Environ Challenges. 2024;14:100979. 10.1016/j.envc.2023.100826

3. Karch PS, Asidi N, Manzambi ZM, Salaun JJ. La faune anophélienne à Kinshasa (Zaïre) et la transmission du paludisme humain. Bull Soc Pathol Exot. 1992;85:123–9. PMID: 1446181

4. Karch PS, Asidi N, Manzambi ZM, Salaun JJ. La faune culicidienne et sa nuisance à Kinshasa (Zaïre). Bull Soc Pathol Exot. 1993;86:231–6. PMID: 8504267

5. Coene J. Malaria in urban and rural Kinshasa: the entomological input. Med Vet Entomol. 1993;7(2):127–37. 10.1111/j.1365-2915.1993.tb00665.x

6. Riveron JM, Watsenga F, Irving H, Irish SR, Wondji CS. High Plasmodium infection rate and reduced bed net efficacy in multiple insecticide-resistant malaria vectors in Kinshasa, Democratic Republic of the Congo. J Infect Dis. 2018;217(2):320–8. 10.1093/infdis/jix570

7. Nguiffo-Nguete D, Mugenzi LMJ, Manzambi EZ, Tchouakui M, Wondji M, Tekoh T, et al. Evidence of intensification of pyrethroid resistance in the major malaria vectors in Kinshasa, Democratic Republic of the Congo. Sci Rep. 2023;13(1):18645. 10.1038/s41598-023-41952-2

8. Wat’senga F, Manzambi EZ, Lunkula A, Mulumbu R, Mampangulu T, Lobo N, et al. Nationwide insecticide resistance status and biting behaviour of malaria vector species in the Democratic Republic of the Congo. Malar J. 2018;17(1):129. 10.1186/s12936-018-2285-6.

9. Kashamuka MM, Banek K, White SJ, Atibu JL, Mvuama NM, Bala JAM, et al. Baseline characteristics for phase II of the Kinshasa malaria cohort study: cohort profile. BMJ Open. 2024;14(11):e085360. 10.1136/bmjopen-2024-085360

10. Ferrari G, Ntuku HM, Schmidlin S, Diboulo E, Tshefu AK, Lengeler C. A malaria risk map of Kinshasa, Democratic Republic of the Congo. Malar J. 2016;15(1):27. 10.1186/s12936-015-1074-8

11. République Démocratique du Congo, Ministère de la Santé Publique, Programme National de Lutte contre le Paludisme. Plan stratégique national 2016–2020. Kinshasa; 2016.

12. U.S. President’s Malaria Initiative. Democratic Republic of the Congo malaria operational plan FY 2023 [Internet]. 2023. Available from: https://www.pmi.gov

13. Gatton ML, Chitnis N, Churcher T, Donnelly MJ, Ghani AC, Godfray HCJ, et al. The importance of mosquito behavioural adaptations to malaria control in Africa. Evolution. 2013;67(4):1218–30. 10.1111/evo.12063

14. Bhatt S, Weiss DJ, Cameron E, Bisanzio D, Mappin B, Dalrymple U, et al. The effect of malaria control on Plasmodium falciparum in Africa between 2000 and 2015. Nature. 2015;526(7572):207–11. 10.1038/nature15535

15. Mwandagalirwa MK, Levitz L, Thwai KL, Parr JB, Goel V, Janko M, et al. Individual and household characteristics of persons with Plasmodium falciparum malaria in sites with varying endemicities in Kinshasa Province, Democratic Republic of the Congo. Malar J. 2017;16(1):456. 10.1186/s12936-018-2429-8

16. Gillies MT, De Meillon B. The Anophelinae of Africa south of the Sahara (Ethiopian Zoogeographical Region). 2nd ed. Johannesburg: South African Institute for Medical Research; 1968. 343 p.

17. Coetzee M. Key to the females of Afrotropical Anopheles mosquitoes (Diptera: Culicidae). Malar J. 2020;19(1):70. 10.1186/s12936-020-3144-9

18. World Health Organization. Test procedures for insecticide resistance monitoring in malaria vector mosquitoes. Geneva: WHO; 2016.

19. Salako AS, Ossè R, Padonou GG, Dagnon F, Aïkpon R, Kpanou C, et al. Population dynamics of Anopheles gambiae s.l. and Culex quinquefasciatus in rural and urban settings before an indoor residual spraying campaign in northern Benin. Vector Borne Zoonotic Dis. 2019;19(9):674–84. 10.1089/vbz.2018.2409

20. Santolamazza F, Mancini E, Simard F, Qi Y, Tu Z, Della Torre A. Insertion polymorphisms of SINE200 retrotransposons within speciation islands of Anopheles gambiae molecular forms. Malar J. 2008;7(1):163. 10.1186/1475-2875-7-163

21. Wirtz RA, Burkot TR, Andre RG, Rosenberg R, Collins WE, Roberts DR. Identification of Plasmodium vivax sporozoites in mosquitoes using an enzyme-linked immunosorbent assay. Am J Trop Med Hyg. 1985;34(6):1048–54. 10.4269/ajtmh.1985.34.1048

22. De Arruda ME, Collins KM, Hochberg LP, Ryan PR, Wirtz RA, Ryan JR. Quantitative determination of sporozoites and circumsporozoite antigen in mosquitoes infected with Plasmodium falciparum or P. vivax. Ann Trop Med Parasitol. 2004;98(2):121–7. 10.1179/000349804225003181

23. Beier JC, Asiago CM, Onyango FK, Koros JK. ELISA absorbance cut-off method affects malaria sporozoite rate determination in wild Afrotropical Anopheles. Med Vet Entomol. 1988;2(3):259–64. 10.1111/j.1365-2915.1988.tb00193.x

24. Huynh LY, Sandve SR, Hannan LM, Van Ert M, Gimnig JE. Fitness costs of pyrethroid insecticide resistance in Anopheles gambiae. Christchurch, New Zealand; 2007.

25. Acford-Palmer H, Campos M, Bandibabone J, N’Do S, Bantuzeko C, Zawadi B, et al. Detection of insecticide resistance markers in Anopheles funestus from the Democratic Republic of the Congo using a targeted amplicon sequencing panel. Sci Rep. 2023;13(1):17363. 10.1038/s41598-023-44457-0

26. Lines JD, Curtis CF, Wilkes TJ, Njunwa KJ. Monitoring human-biting mosquitoes (Diptera: Culicidae) in Tanzania with light traps hung beside mosquito nets. Bull Entomol Res. 1991;81(1):77–84. 10.1017/S0007485300053268

27. Namango IH, Marshall C, Saddler A, Ross A, Kaftan D, Tenywa F, et al. The CDC light trap and the human decoy trap compared to the human landing catch for measuring Anopheles biting in rural Tanzania. Malar J. 2022;21(1):181. 10.1186/s12936-022-04192-9

28. Overgaard HJ, Sæbø S, Reddy MR, Reddy VP, Abaga S, Matias A, et al. Light traps fail to estimate reliable malaria mosquito biting rates on Bioko Island, Equatorial Guinea. Malar J. 2012;11(1):56. 10.1186/1475-2875-11-56

29. Magbity EB, Lines JD, Marbiah MT, David K, Peterson E. How reliable are light traps in estimating biting rates of adult Anopheles gambiae s.l. (Diptera: Culicidae) in the presence of treated bed nets? Bull Entomol Res. 2002;92(1):71–6. 10.1079/BER2001131

30. Briët OJT, Huho BJ, Gimnig JE, Bayoh N, Seyoum A, Sikaala CH, et al. Applications and limitations of CDC miniature light traps for measuring biting densities of African malaria vector populations: a pooled analysis of 13 comparisons with human landing catches. Malar J. 2015;14(1):247. 10.1186/s12936-015-0761-9

31. Dida GO, Anyona DN, Abuom PO, Akoko D, Adoka SO, Matano AS, et al. Spatial distribution and habitat characterization of mosquito species during the dry season along the Mara River and its tributaries in Kenya and Tanzania. Infect Dis Poverty. 2018;7(1):2. 10.1186/s40249-017-0385-0

32. Metelo E, Bukaka E, Luemba TB, Situakibanza H, Sangaré I, Mesia G, et al. Détermination des paramètres bioécologiques et entomologiques d’Anopheles gambiae s.l. dans la transmission du paludisme à Bandundu-ville, République Démocratique du Congo. Pan Afr Med J. 2015;22:1–10. 10.11604/pamj.2015.22.108.6774

33. Kahamba NF, Tarimo FS, Kifungo K, Mponzi W, Kinunda SA, Simfukwe A, et al. Societal uses of main water bodies inhabited by malaria vectors and implications for larval source management. Malar J. 2024;23(1):336. 10.1186/s12936-024-05154-z

34. Akuoko OK, Dhikrullahi SB, Hinne IA, Mohammed AR, Owusu-Asenso CM, Coleman S, et al. Biting behaviour, spatio-temporal dynamics, and insecticide resistance status of malaria vectors in different ecological zones in Ghana. Parasit Vectors. 2024;17(1):16. 10.1186/s13071-023-06065-9

35. Zanga J, Metelo E, Mvuama N, Nsabatien V, Mvudi V, Banzulu D, et al. Species composition and distribution of the Anopheles gambiae complex circulating in Kinshasa. GigaByte. 2024; 2023:1–12. 10.46471/gigabyte.104

36. Sinka ME, Bangs MJ, Manguin S, Coetzee M, Mbogo CM, Hemingway J, et al. The dominant Anopheles vectors of human malaria in Africa, Europe and the Middle East: occurrence data, distribution maps and bionomic précis. Parasit Vectors. 2010;3(1):117. 10.1186/1756-3305-3-117

37. Zanga J, Metelo E, Mbanzulu K, Irish S, Mulenda B, Di-Mosi Wumba R, et al. Susceptibility status of Anopheles gambiae s.l. to insecticides used for malaria control in Kinshasa, Democratic Republic of the Congo. Ann Afr Med. 2022;15(2):4533. Available from: 10.4314/aam.v15i2.2

38. Metelo E, Zanga J, Batumbo D, Mandja BA, Lukoki H, Bokulu A, et al. Complexity of vector control and entomological surveillance in endemic sentinel sites of the National Malaria Control Program (NMCP) in the Democratic Republic of the Congo (DRC) [Internet]. 2024. Available from: https://www.intechopen.com

39. Sherrard-Smith E, Skarp JE, Beale AD, Fornadel C, Norris LC, Moore SJ, et al. Mosquito feeding behavior and how it influences residual malaria transmission across Africa. Proc Natl Acad Sci U S A. 2019;116(30):15086–95. 10.1073/pnas.1820646116

40. Sougoufara S, Ottih EC, Tripet F. The need for new vector control approaches targeting outdoor biting anopheline malaria vector communities. Parasit Vectors. 2020;13(1):295. 10.1186/s13071-020-04170-7

41. Githeko AK, Adungo NI, Karanja DM, Hawley WA, Vulule JM, Seroney IK, et al. Some observations on the biting behavior of Anopheles gambiae s.s., Anopheles arabiensis, and Anopheles funestus and their implications for malaria control. Exp Parasitol. 1996;82(3):306–15. 10.1006/expr.1996.0038

42. Bobanga T, Umesumbu SE, Mandoko AS, Nsibu CN, Dotson EB, Beach RF, et al. Presence of species within the Anopheles gambiae complex in the Democratic Republic of the Congo. Trans R Soc Trop Med Hyg. 2016;110(6):373–5. 10.1093/trstmh/trw035

43. Sharma A, Kinney NA, Timoshevskiy VA, Sharakhova MV, Sharakhov IV. Structural variation of the X chromosome heterochromatin in the Anopheles gambiae complex. Genes (Basel). 2020;11(3):327. 10.3390/genes11030327

44. Bobanga T, Ayieko W, Zanga M, Umesumbu S, Landela A, Fataki O, et al. Field efficacy and acceptability of PermaNet® 3.0 and OlysetNet® in Kinshasa, Democratic Republic of the Congo. J Vector Borne Dis. 2013;50(3):206–14. PMID: 24220080

45. Wat’senga F, Agossa F, Manzambi EZ, Illombe G, Mapangulu T, Muyembe T, et al. Intensity of pyrethroid resistance in Anopheles gambiae before and after a mass distribution of insecticide-treated nets in Kinshasa and in 11 provinces of the Democratic Republic of Congo. Malar J. 2020;19(1):169. 10.1186/s12936-020-03240-6

46. Metelo-Matubi E, Zanga J, Binene G, Mvuama N, Ngamukie S, Nkey J, et al. The effect of a mass distribution of insecticide-treated nets on insecticide resistance and entomological inoculation rates of Anopheles gambiae s.l. in Bandundu City, Democratic Republic of the Congo. Pan Afr Med J. 2021;40:1–10. 10.11604/pamj.2021.40.118.27365

47. De Marco CM, Virgillito C, Frosi L, Santarelli G, Filipponi F, Manica M, et al. Habitat drivers and predicted distribution shifts of Anopheles coluzzii under climate change: results from a systematic review. Sci Total Environ. 2025;992:179939. 10.1016/j.scitotenv.2025.179939

48. Longo-Pendy NM, Tene-Fossog B, Tawedi RE, Akone-Ella O, Toty C, Rahola N, et al. Ecological plasticity to ion concentration determines genetic response and dominance of Anopheles coluzzii larvae in urban coastal habitats of Central Africa. Sci Rep. 2021;11(1):15781. 10.1038/s41598-021-94258-6

49. Costantini C, Ayala D, Guelbeogo WM, Pombi M, Some CY, Bassole IH, et al. Living at the edge: biogeographic patterns of habitat segregation conform to speciation by niche expansion in Anopheles gambiae. BMC Ecol. 2009;9(1):16. 10.1186/1472-6785-9-16

50. Diabaté A, Dabiré RK, Heidenberger K, Crawford J, Lamp WO, Culler LE, et al. Evidence for divergent selection between the molecular forms of Anopheles gambiae: role of predation. BMC Evol Biol. 2008;8(1):5. 10.1186/1471-2148-8-5

51. Diabaté A, Dabiré RK, Kim EH, Dalton R, Millogo N, Baldet T, et al. Larval development of the molecular forms of Anopheles gambiae (Diptera: Culicidae) in different habitats: a transplantation experiment. J Med Entomol. 2005;42(4):548–53. 10.1093/jmedent/42.4.548

52. Carrel M, Kim S, Mwandagalirwa MK, Mvuama N, Bala JA, Nkalani M, et al. Individual, household and neighborhood risk factors for malaria in the Democratic Republic of the Congo support new approaches to programmatic intervention. Health Place. 2021;70:102581. 10.1016/j.healthplace.2021.102581

53. Sendor R, Banek K, Kashamuka MM, Mvuama N, Bala JA, Nkalani M, et al. Epidemiology of Plasmodium malariae and Plasmodium ovale spp. in a highly malaria-endemic country: a longitudinal cohort study in Kinshasa Province, Democratic Republic of the Congo. Malar J. 2023;22(1):97. 10.1101/2023.04.20.23288826

54. Diallo AO, Banek K, Kashamuka MM, Bala JAM, Nkalani M, Kihuma G, et al. Impact of malaria diagnostic choice on monitoring Plasmodium falciparum prevalence estimates in the Democratic Republic of the Congo and relevance to control programs in high-burden countries. PLOS Glob Public Health. 2023;3(7):e0001375. 10.1371/journal.pgph.0001375

